# Body size mediates latitudinal population differences in response to *Bd* infection in two amphibian species

**DOI:** 10.1101/2021.07.16.452656

**Authors:** Sara Meurling, Maria Cortazar-Chinarro, Mattias Siljestam, David Åhlen, Erik Ågren, Jacob Höglund, Anssi Laurila

## Abstract

Populations of the same species may differ in their sensitivity to pathogens but the factors behind this variation are poorly understood. Moreover, infections may cause sub-lethal fitness effects even in species resistant or tolerant to disease. The chytrid fungus *Batrachochytrium dendrobatidis* (*Bd*), is a generalist pathogen which has caused amphibian population declines worldwide. In many species, *Bd* infection causes the disease chytridiomycosis, often leading to high mortality. We investigated how geographical origin affects tolerance to *Bd* by exposing newly metamorphosed individuals of two North European amphibians (moor frog *Rana arvalis*, common toad *Bufo bufo*) from two latitudinal regions to two different *Bd*GPL strains. *Bd* exposure strongly lowered survival in *B. bufo*, and in both species survival was lower in the northern region, this difference being much stronger in *B. bufo*. Northern individuals were smaller in both species, and the survival difference between the regions was size-mediated with smaller individuals being more sensitive to *Bd*. In both species, *Bd* exposure led to sub-lethal effects in terms of reduced growth suggesting that even individuals surviving the infection may have reduced fitness mediated by smaller body size. *Bd* strain affected size-dependent mortality differently in the two regions. We discuss the possible mechanisms how body size and geographical origin can contribute to the present results.

## Introduction

Natural populations are increasingly affected by emerging infectious diseases (Daszak et al. 2000, Pennisi 2010, Fisher et al. 2012, Scheele et al. 2019). Many of the emerging diseases are caused by generalist and opportunistic fungal pathogens which can infect a wide range of host species (Wibbelt et al. 2010, Fisher et al. 2012, Lorch et al. 2016, More et al. 2018). The virulence of a fungal pathogen often differs among host species, leading to population declines in some hosts while having no apparent effect on others (Casadevall 2007, Herceg et al. 2021). There is also evidence that populations of a host species may differ in their susceptibility, but apart from plant systems few studies have addressed this question in detail (Ebert 2008; Laine et al. 2011; Bradley et al. 2015; Martin-Torrijos et al. 2017).

The chytrid fungus *Batrachochytrium dendrobatidis* (*Bd*), causing the disease chytridiomycosis in amphibians, is a generalist pathogen which has caused the decline of over 500 amphibian species, including the presumed extinction of 90 species (Berger et al. 1998, Skerratt et al. 2007, Lips 2016, Scheele et al. 2019). *Bd* is endemic in East Asia where itcoexists with the native fauna, but severe outbreaks of chytridiomycosis have been observed in the Americas and Australia (Lips 2016, Scheele et al. 2019). There is considerable variation in virulence among genetic strains of *Bd* (Farrer et al. 2011, Bataille et al. 2013, Greenspan et al. 2018) and *Bd*GPL, the global panzootic lineage originating in Eastern Asia, has caused most of the chytridiomycosis outbreaks (O’Hanlon et al. 2018). While genetic variation within *Bd*GPL is relatively limited (O’Hanlon et al. 2018), there is evidence for virulence differences between *Bd*GPL strains (Becker et al. 2017; Burrow et al. 2017, Dang et al 2017). As a generalist pathogen, *Bd* infects a wide range of amphibian species (Fisher et al. 2012, Olson et al. 2013). However, all species do not develop chytridiomycosis; many are resistant to the disease and can clear the infection, while others can tolerate high infection loads without developing the disease (Fisher et al. 2009, Gahl et al. 2012, Ellison et al. 2014, Scheele et al. 2017). Similarly, geographical populations of the same species can differ in their susceptibility to *Bd* (Savage and Zamudio 2011, Bradley et al. 2015, Kosch et al. 2019). These differences can be due to genetic differences in traits like immune response and behavior (Richards-Zawacki 2010), and are in some cases linked with direct *Bd*-mediated selection (Savage & Zamudio 2016, Savage et al. 2018). Although infection does not cause direct mortality in the resistant and tolerant species, sub-lethal fitness effects such as decreased growth have been detected (Bielby et al. 2015, Burrow et al. 2017, Campbell et al. 2019).

Climate-related latitudinal divergence is an important structuring force of intraspecific genetic variation (e.g., Hewitt 2000, Conover et al. 2009), but its potential effects mediating host-pathogen interactions have received little attention. Two lines of evidence suggest that amphibian populations living at high latitudes in the northern hemisphere may be especially vulnerable to disease. Firstly, due to post-glacial colonization patterns northern populations often harbor less genetic variation (Hewitt 2000). In many amphibians, this is true also for immunogenetic variation in major histocompatibility (MHC) genes (Zeisset and Beebee 2014, Cortázar-Chinarro et al. 2017), which is associated with *Bd* resistance (Savage and Zamudio 2011, Savage et al. 2018, Kosch et al. 2019). Furthermore, pathogen richness and abundance are significant predictors of adaptive MHC variation (Wang et al. 2017). As pathogen richness and abundance decrease towards colder climates (Schemske et al. 2009), populations at higher latitudes may encounter lower diversity and a lower number number of pathogens which may lead to increased drift and loss of adaptive immunogenetic variation in these populations (Cortázar-Chinarro et al. 2017). Secondly, time-constrained high-latitude environments select for high larval growth and development rates (Palo et al. 2003, Luquet et al. 2019), which in amphibians can trade-off with disease resistance (Johnson et al. 2011, Woodhams et al. 2016) and immune response (Gervasi and Foufopoulos 2007, Murillo-Rincon et al. 2017). While all these factors may contribute to lower ability to withstand novel pathogens in high-latitude populations, no studies on disease resistance between latitudinal populations have been made. Here we conducted a laboratory common garden experiment to examine inter- and intraspecific population differences in response to *Bd* infection. Our aims were three-fold: 1) to investigate the response of two common north European amphibians (moor frog *Rana arvalis* and common toad *Bufo bufo*) to *Bd* infection, 2) investigate if the responses differ between southern and northern Scandinavian populations of these species and 3) evaluate if these responses differ between two geographically separated *Bd* lineages. To this end, we infected newly metamorphosed amphibians with two different *Bd*GPL lineages and measured their survival and growth during a 30-day exposure period.

## Methods

### Animal rearing

Both *R. arvalis* (hereafter *Ra*) and *B. bufo* (hereafter *Bb*) are widespread amphibians in Europe occurring up to the polar circle in the north (Sillero et al. 2014). Both species are explosive breeders and mate in early spring. In southern Sweden, *Bd* prevalence in breeding adults is 15.3% (n = 288) and 3.4% (n = 941) in *Ra* and *Bb*, respectively (Meurling et al. 2020).

Eggs of both species were collected in April 2016 at two sites in Skåne county in southernmost Sweden and May 2016 at two sites in in Norrbotten county in northern Sweden (Fig. 1; Table S1). We collected approximately ten eggs from each of ten different clutches at each site. The eggs and tadpoles were reared in walk-in climate-controlled rooms at Uppsala University in plastic tanks filled with 20l reconstituted soft water (RSW; NaHCO_3_, CaSO_4_, MgSO_4_ and KCl added to deionized water; APHA 1985) until metamorphosis. Each clutch was kept in a separate tank under 18:6 h light/dark regime at 19°C. The tadpoles were fed *ad libitum* spinach and fish flakes and water was changed every third day. At metamorphic climax (stage 42; Gosner 1960), the animals were moved to another tank of the same size with access to aquatic and terrestrial (aquarium sand) habitat and a shelter. Four days after completion of tail absorption (stage 46), the animals were transported to the sealed experimental facilities at the Swedish Institute for Veterinary Science, Uppsala, where they were kept individually in 1.2 l plastic tanks lined with moist paper towels and a lid of a plastic bottle as a shelter. The metamorphs were kept in these tanks until the end of the experiment and fed fruit flies and crickets *ad libitum* under 18/6h light/dark regime at 19°C. The condition of each animal was checked daily and the tanks were cleaned every third day.

**Fig. 1.**
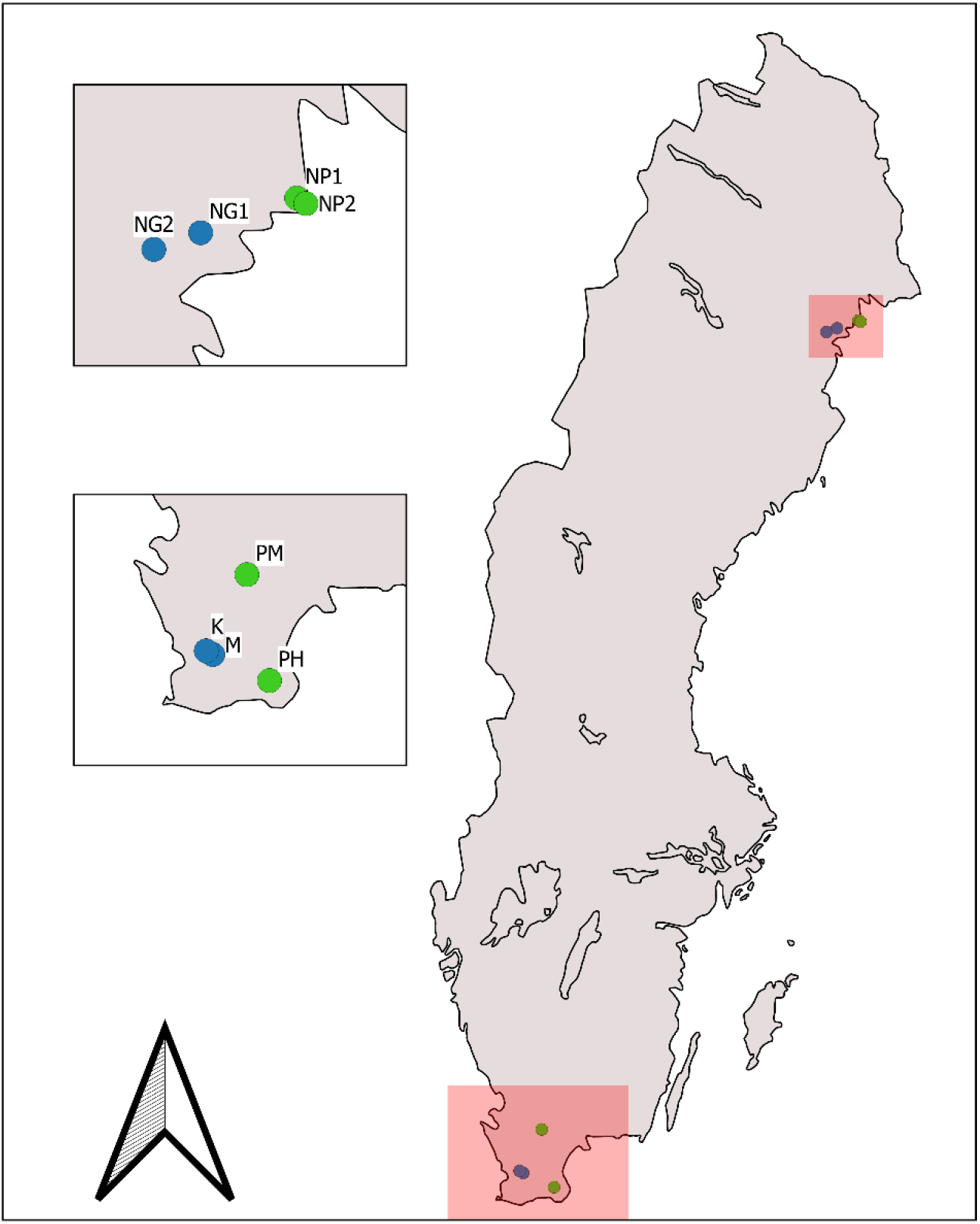
Map of Sweden showing the sites of egg collection. Blue and green circles show *R. arvalis* and green *B. bufo* sites, respectively.

### Infection experiment

The infection treatments were conducted after one week of acclimatization at the experimental facility. The experimental animals were exposed to one of two isolates of *Bd*-GPL (SWE or UK) or a sham infection consisting of culture medium (Table 1). The UK isolate (UKMal 01) was isolated from a wild alpine newt (*Ichthyosaura alpestris*) in the UK in 2008. The Swedish isolate (SWED-40-5) originated from a wild green toad (*Bufotes viridis*) in Malmö municipality in southern Sweden in 2015. The animals were exposed individually for 5 h to 200μl culture media containing a dosage of 60 000 zoospores from one of the *Bd* strains in 30 ml of RSW. The control group (C) was exposed for 5 h to an equivalent volume of RSW and culture media without *Bd* spores. Altogether, we treated 74 (25 in SWE, 24 in UK and 25 in C treatment) southern and 46 (16 SWE, 14 UK, 16 C) northern *Ra*. The corresponding numbers for *Bb* were 64 (21, 19, 24) southern and 90 (31, 31, 28) northern individuals.

**Table 1.** Number of experimental and infected individuals (as determined by qPCR) in different infection treatments.

| Species | Region | Treatment | Total<br>Nb. | Positive | % | Negative | % | Failed | % |
| --- | --- | --- | --- | --- | --- | --- | --- | --- | --- |
| *Bufo bufo* | North | Control | 28 | 1 | 3,6 | 16 | 57,1 | 11 | 39,3 |
|  |  | *BdSWE* | 31 | 31 | 100,0 | 0 | 0,0 | 3 | 9,7 |
|  |  | *BdUK* | 31 | 31 | 100,0 | 0 | 0,0 | 0 | 0,0 |
|  | South | Control | 24 | 9 | 37,5 | 13 | 54,2 | 2 | 8,3 |
|  |  | *BdSWE* | 21 | 21 | 100,0 | 0 | 0,0 | 3 | 14,3 |
|  |  | *BdUK* | 19 | 14 | 73,7 | 1 | 5,3 | 4 | 21,1 |
| *Rana arvalis* | North | Control | 16 | 1 | 6,3 | 6 | 37,5 | 9 | 56,3 |
|  |  | *BdSWE* | 16 | 16 | 100,0 | 0 | 0,0 | 1 | 6,3 |
|  |  | *BdUK* | 14 | 13 | 92,9 | 0 | 0,0 | 1 | 7,1 |
|  | South | Control | 25 | 2 | 8,0 | 18 | 72,0 | 5 | 20,0 |
|  |  | *BdSWE* | 25 | 25 | 100,0 | 0 | 0,0 | 0 | 0,0 |
|  |  | *BdUK* | 24 | 23 | 95,8 | 1 | 4,2 | 0 | 0,0 |

After exposure the animals were monitored for 30 days, or until death. Animals showing irreversible signs of chytridiomycosis (loss of righting function) were euthanized with an overdose of MS222. Body mass of the animals was measured at the start and end of the experiment (or at death). At the end of the experiment, the surviving animals were euthanized and stored in 96% ethanol at 4 °C.

### DNA extraction and qPCR analyses

To confirm infection status, we assessed the presence of *Bd* by using qPCR. DNA was extracted from a hind leg using a Prepman Ultra method described in Boyle et al. (2004). Presence of *Bd* was assessed by amplifying the internal transcribed spacer (ITS)-5.8S rRNA region (Boyle et al 2004). 25μl reactions containing 12.5μl 2X Taqman Master Mix (Applied Biosystem, ref. 4318157), 2.25 μl 10μM each of forward and reverse primers, 0.625 μl 10μM MGB probe and 5μl of DNA (diluted x10 in water) were run. Each sample was run in triplicate. An exogenous internal positive control (IPC; (Hyatt et al. 2007)) was added to one well in each triplicate (1μl 10XExo IPC master mix and 0.5μl 50XExo IPC DNA to each sample) (VICTIM dye, Applied Biosystems ref. 4304662) to avoid false negatives due to inhibitors. The qPCR assays were run on a Biorad CFX96 Real Time System machine using amplification conditions described in Boyle et al. (2004) with standards of 0.1, 1, 10 and 100 genomic equivalents (GE). An individual was recorded as positive if at least one of the triplicate samples exhibited a positive signal (i.e. an exponential amplification curve). If the IPC showed signs of inhibition, negative samples were rerun once before the samples were assigned as not scoreable (NA) and removed from the data set. The above-mentioned standards were used to create a standard curve which was then used to calculate the infection intensity for each individual expressed in genome equivalents (GE).

For the statistical analysis of the infection, we used the logarithm (base 10, zero values were replaced by 0.001, one tenth of the lowest measured non-zero value) of GE and refer to this as infection load (IL).

### Statistical analyses

All analyses were conducted in R 3.5.2 (R Core Team 2018). Survival was analysed using Generalised Linear Models with a binomial error distribution and a logit link function while data on growth and IL were analysed using linear models. The model assumptions of the linear models were checked using the model diagnostic plots in R. In *Bb*, the growth data were log-transformed due to heteroscedasticity. Consequently, growth in was defined as proportional growth per day: log(mass at death /mass at exposure)/lifespan) for both species.

The models that best explained differences in IL, survival and growth were selected using bidirectional elimination (*stepAIC* function in R package *MASS*) starting from the full model with Response ~ Region + *Bd*-strain + Size at infection + Infection load + Interactions. For IL, the full model was IL ~ Region + *Bd*-strain + Size at infection + Interactions. The two populations within each region were pooled in the analyses as our interest was increasing the sampled genetic variation within each region rather than studying population differences, and our experimental design did not allow for effective tests of population effects.

To separate the effect of each *Bd*-treatment we also ran the models without the control treatment following the same general structure. Differences in size at infection between regions and treatment were analysed with factorial ANOVAs. This was also done for differences in IL between the species.

## Results

The qPCR analyses showed high infection success: only two of the successfully analysed individuals from the exposed groups were negative to *Bd* infection at the end of the experiment (one *Bb* and one *Ra*; Table 1) and were removed from the analyses. Thirteen control individuals were also found *Bd* positive (Table 1). However, the infection intensity was very low and we find it likely that these samples were contaminated during sample processing at the end of the experiment. For the statistical analysis, their infection intensity was therefore considered to be 0 GE.

At the start of the experiment, northern animals were significantly smaller than southern animals both in *Ra* (*F*_1, 115_ = 25.05, p < 0.001) and in *Bb* (*F*_1, 149_ = 156.94 p < 0.001; Fig. 2). There was no difference in size at exposure between treatments (*Ra*: *F*_2, 115_ = 0.15, p = 0.86, *Bb*: *F*_2, 149_ = 0.9, p = 0.41).

**Fig. 2.**
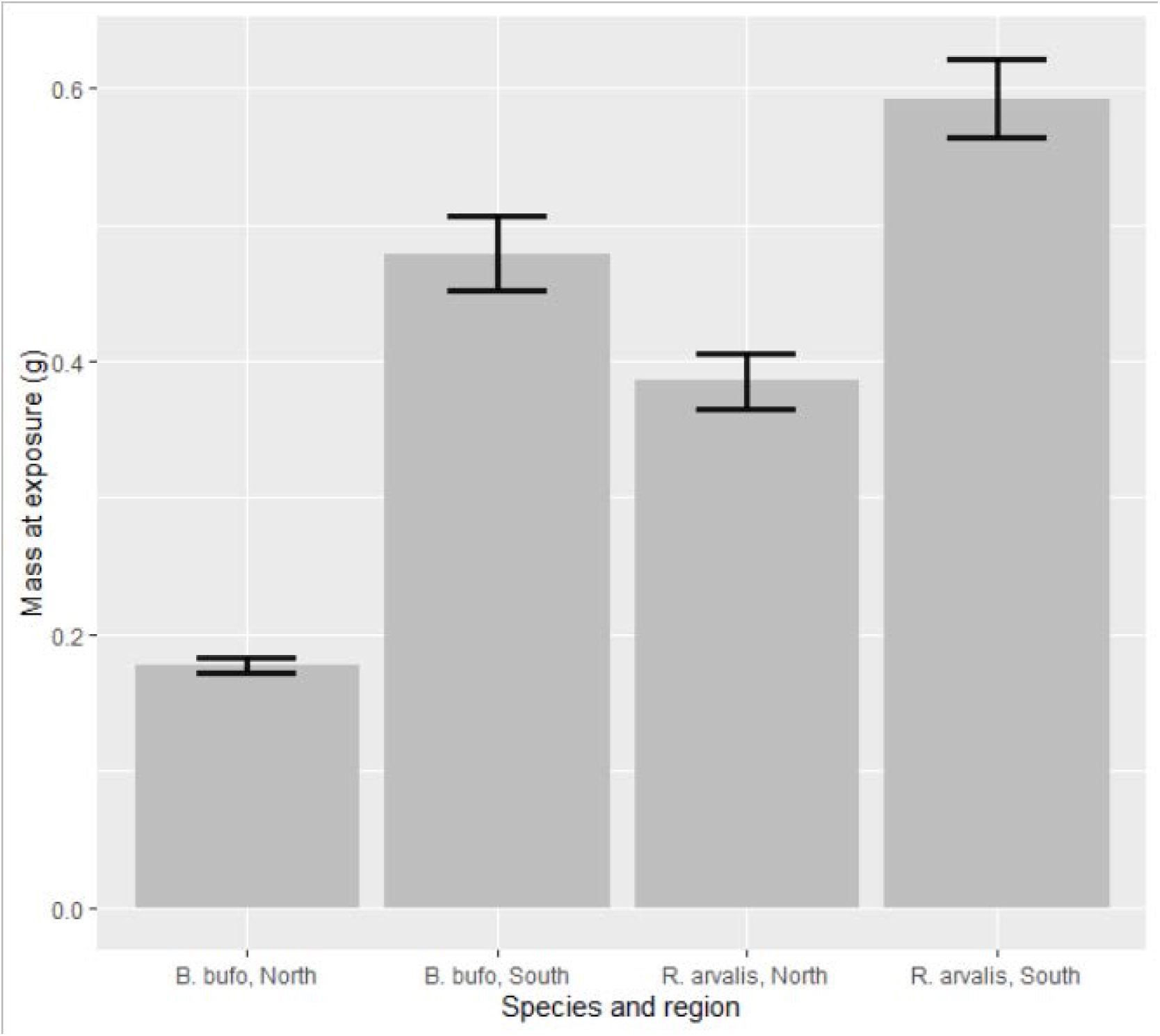
Mass at exposure (g ± SE) per region for *R. arvalis* and *B. bufo*

### Infection load

IL differed between species (*F*_1, 163_ = 15.09, p < 0.001, Fig S1), *Bb* having higher loads. The selected model for *Ra* was IL ~ Size + Region + Size × Region (Table S2a). We found a significant effect of the interaction between size and region on IL (*F*_1, 69_ = 6.5, p = 0.013)., Size at infection had a negative effect on IL in the northern region, while in the southern region size had no effect (Fig. 3a).

**Fig. 3.**
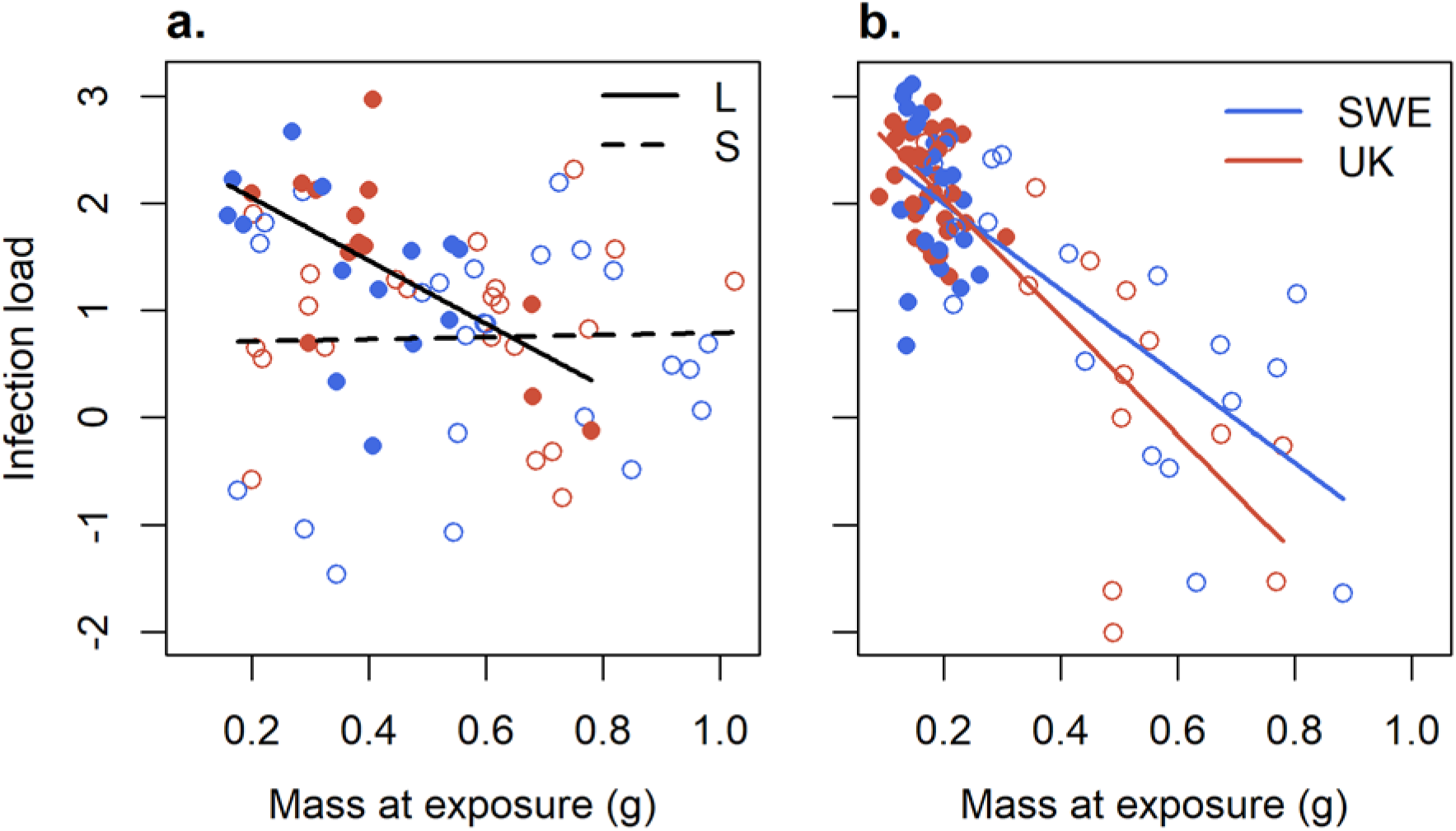
Infection load (log10 average GE, at the end of treatment) as the function of size at infection with either the Swedish or UK strain in **a.** *R. arvalis* and **b.** *B. bufo*. Lines give the predictions of the model. Filled dots and solid lines represent the northern region, while open dots and dashed lines represent the southern region.

For *Bb*, the selected model was IL ~ *Bd*-strain + Size + *Bd*-strain × Size (Table S2b). Size had a significant negative effect on infection load (*F*_1, 87_ = 64.45, p < 0.001, Fig. 3b). The interaction between *Bd*-strain and size was close to significant (*F*_1, 87_ = 3.66, p = 0.059), large toadlets infected with SWE strain having somewhat higher loads than large individuals infected with UK strain.

### Survival

All southern *Ra* survived the experiment, whereas three infected individuals from the northern region died during the experiment, resulting in survival of 92.9 % in the UK and 87.5 % in the SWE treatment. Survival was complete in the control treatment and in the southern region, and these were excluded from the model. The selected model was: Survival ~ Size (Table S3a) indicating poorer survival of smaller individuals in the two infection treatments in the northern region (χ^2^ _1, 26_ = 6.49 p= 0.011; Fig. 4a).

**Fig. 4.**
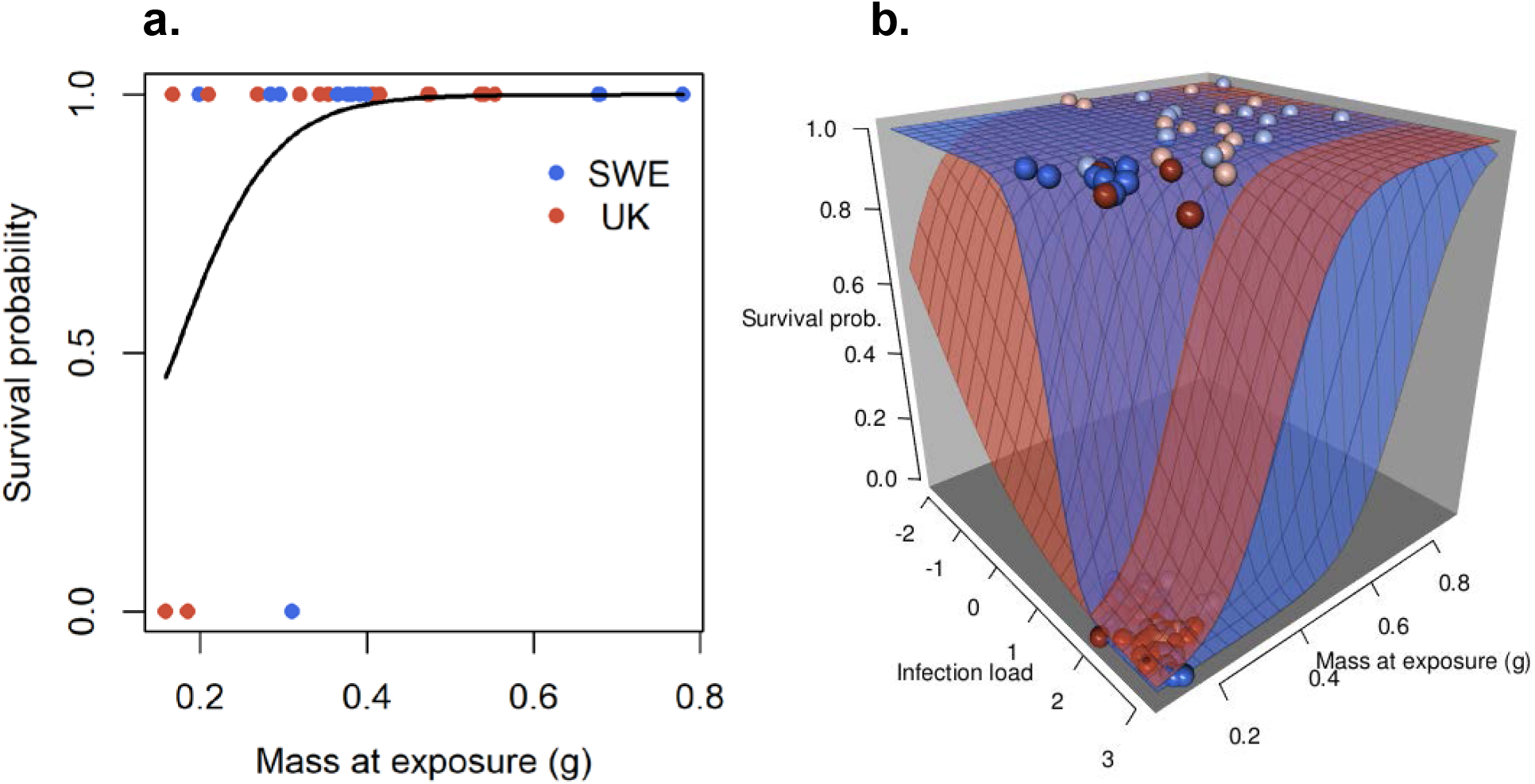
Survival as a function of size at infection with either the Swedish or UK strain for **a**: *R. arvalis* from the northern region. **b**: Survival in *B. bufo* as function of infection load and size at infection, where dark and pale dots represent the northern and southern regions, respectively. Line (**a**) and surfaces (**b**) give the predictions of the model. Blue dots refer to the Swedish *Bd* strain and red dots to the UK strain. In **b** some pale dots are hidden among the dark dots in the lower corner in the front (high infection load and low size).

While all *Bb* in the control treatment survived the experiment, there was considerable mortality in the infection treatments. Furthermore, survival was higher in southern (66.7 % in SWE and 89.5 % in UK) than in the northern region (38.7 % in SWE and 12.9 % in UK). Due to 100% survival in the control treatment we excluded it from the full model. We also excluded the interaction term between region and size as well as region and infection load, as the range of infection load and size from the northern region only covered a small subset of the range of the southern region, and including these interactions may lead to problematic extrapolations. Survival of *Bb* in the two infection treatments was best explained by the model Survival ~ Bd-strain + Size + IL + *Bd*-strain × IL (Table S3b). Initial size had a strong positive effect on survival (*F*_1, 86_ = 13.10, p < 0.001, Fig. 4b), whereas IL had a strong negative effect (*F*_1, 86_ = 27.61, p < 0.001). In addition, the significant interaction between *Bd* strain and infection load (*F*_1, 86_ = 7.05, p = 0.009) corresponded to SWE strain having a steeper slope between survival and infection load compared to UK strain. This results in higher mortality at high and lower mortality at low IL in SWE treatment (Fig. 4b).

### Growth

Growth in *Ra* was best explained by the model Growth ~ Bd-strain + Region + Size + IL + Region × Size + Size × IL (Table S4a). IL had a negative effect on growth in both infected groups (*F*_1, 107_ = 6.16, p = 0.015; Fig. 5a). Size at infection had a negative effect on growth (*F*_1, 107_ = 6.16, p = 0.015, Fig. 5b), the Size × IL interaction (*F*_1, 107_ = 7.21, p = 0.008) was caused by the stronger of IL on growth of small individuals. We also found a significant effect of *Bd* strain on growth (*F*_2, 107_ = 3.13, p = 0.048) and an analysis using only the infected individuals showed that froglets infected with the Swedish strain had an overall lower growth than those infected with the UK strain (*F*_1, 70_ = 5.63, p = 0.02; Fig. 5b).

**Fig. 5.**
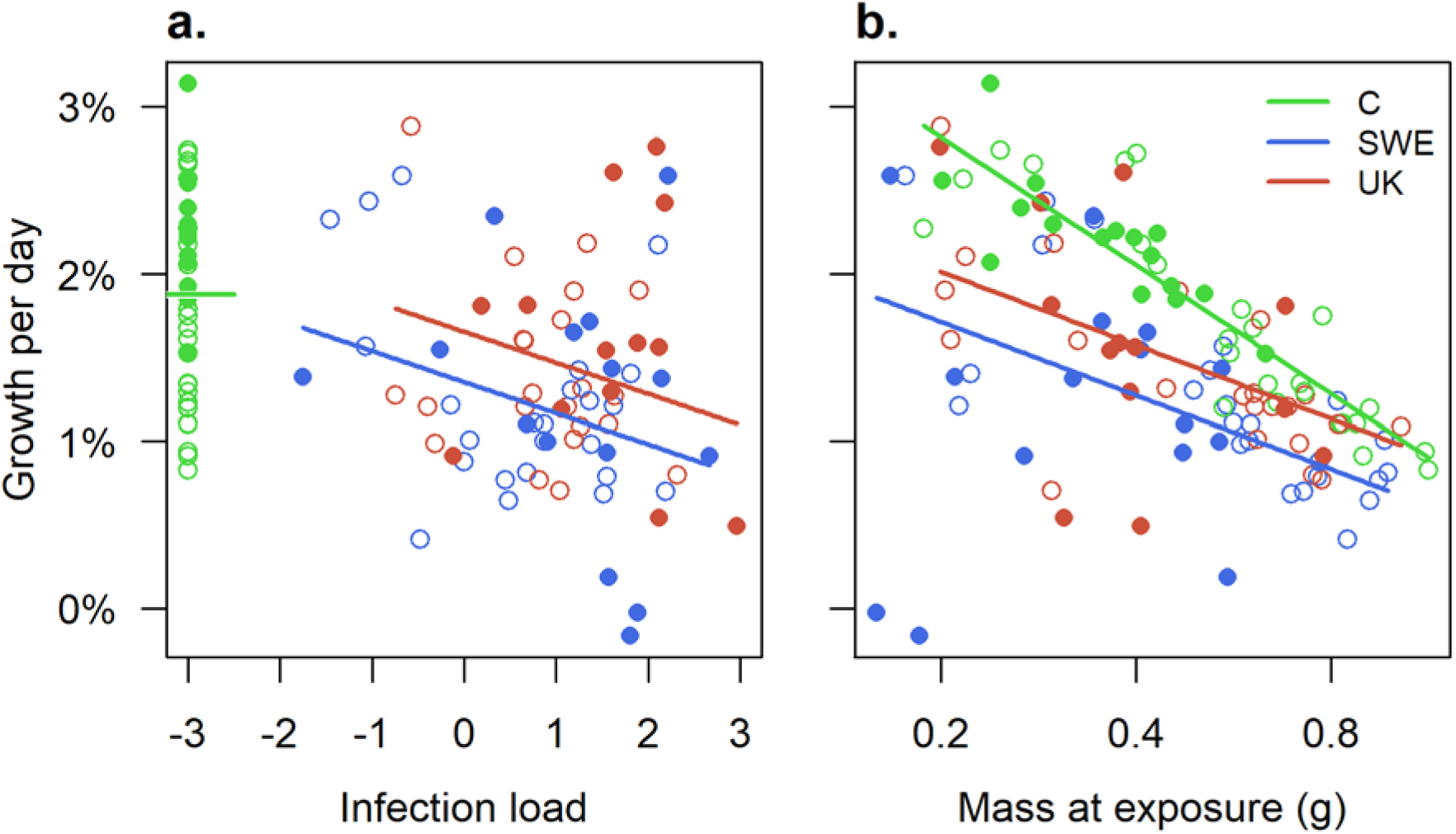
Growth per day (from infection to death or end of experiment) as a function of **a:** infection load (log10 average GE, at the end of treatment) and, **b:** size (at start of treatment) in different infection treatments for *R. arvalis*. Filled and open dots represent the northern and southern regions, respectively. Lines gives the predictions of the model and are evaluated for average infection load in **a** and average size in **b**. Separate models were run for the control group and infected individuals.

Growth analyses in *Bb* revealed strong negative effects of infection load and a size × IL interaction when all three *Bd*-treatments were analysed together (Table S4b). To get more insight on how *Bd* strain affected *Bb* growth we analysed the in the two Bd- infection treatments. This best model was Growth ~ Bd-strain + Region + Size + IL + Bd-strain × Region + Region × Size + Size × IL (Table S4c). The effect of size depended on the region (Region × Size: *F*_1, 83_ = 17.61, p < 0.001) and IL (Size × IL: *F*_1, 83_ = 9.00, p = 0.004). Therefore, we present the effect of size on growth separately for the less and more infected half of the individuals (Fig 6a, b). In a similar manner, how infection load affects growth is presented separately for smaller and larger half of the individuals (Fig 6c, d).

**Fig. 6.**
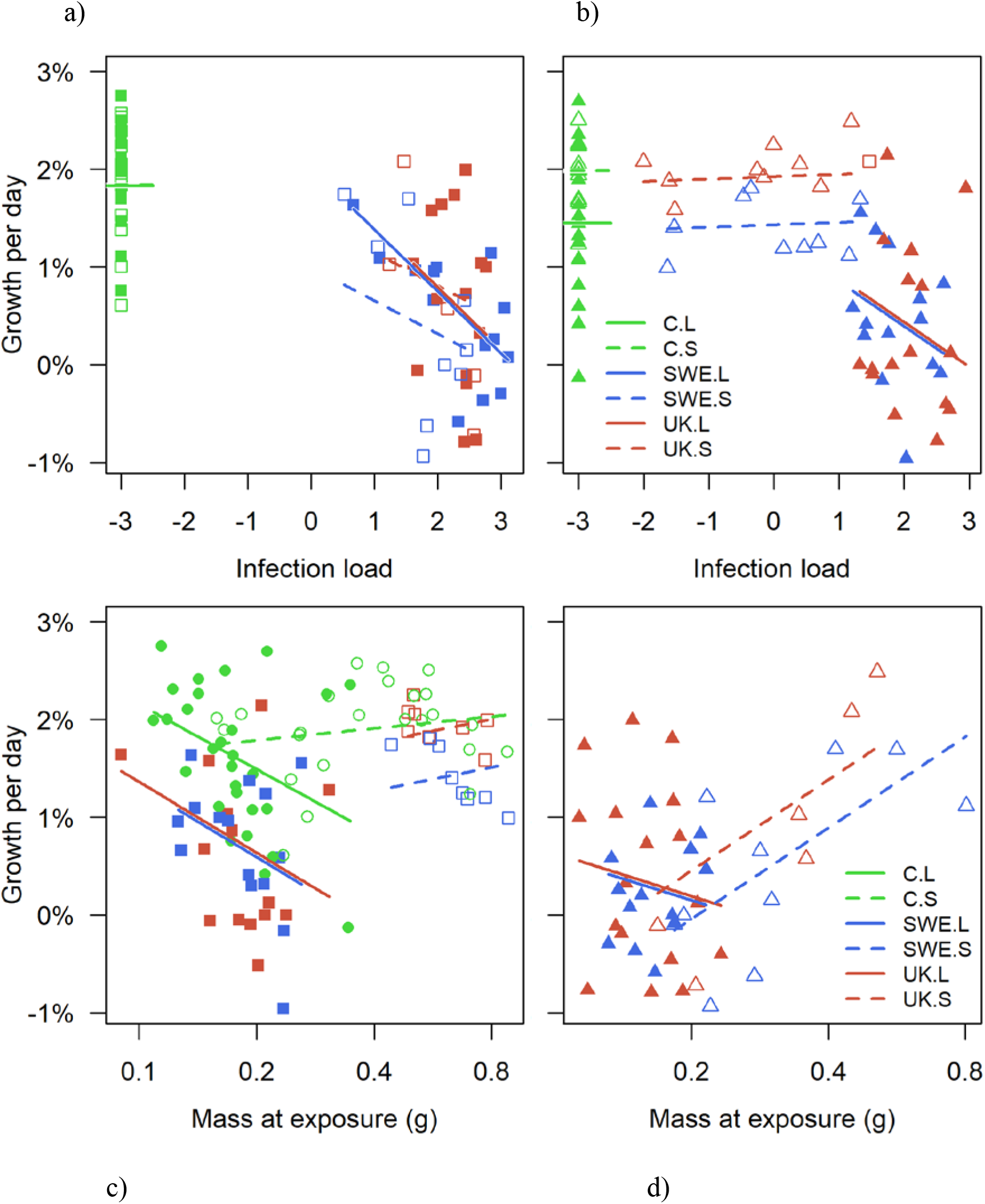
Growth per day as a function of **a-b:** infection load (log10 average GE, at the end of treatment) and, **c-d:** size at infection with either the Swedish or UK strain or no infection (control) for *B. bufo*. Smaller individuals are represented by squares in **a** (smaller than the median of their respective region) and the larger individuals as triangles in **b** (larger or equal to the median of their respective region). The same holds for **c-d** but with respect to infection load. Lines shows model prediction. Filled markers and solid lines represent the northern and open markers and dashed lines represent southern region. Separate models were run for control treated and infected individuals.

In small individuals, IL had strong negative effect on growth in both regions (Fig. 6a). In larger toadlets, growth of southern individuals was unaffected by IL whereas larger northern individuals were negatively affected by IL (Fig. 6b). This was also reflected on how initial mass affected growth: in southern toadlets larger individuals grew proportionally faster, whereas in northern toadlets proportional growth was negatively correlated with initial size (Fig. 6c, d). This difference was especially clear in individuals with higher IL (Fig. 6b). There was no significant difference between the two *Bd*-strains (*F*_1, 83_ = 0.06, p = 0.812) nor significant interaction (*Bd*-strain × Region: *F*_1, 83_ = 2.20, p = 0.142).

## Discussion

We found that *Bd* infection lowered survival in both *Ra* and *Bb*, but the effect of *Bd* was more severe on the latter species, especially in the northern region. Our analyses suggest that the survival differences between the regions were largely mediated by body size, smaller individuals being more sensitive to *Bd*. Furthermore, we found that *Bd* infection led to sub-lethal effects in terms of reduced growth, suggesting that individuals surviving the infection may have lower fitness mediated by their smaller body size. These results suggest that *Bd* infection may have both direct and indirect effects on amphibian populations and that. high latitude populations may run a higher risk of negative effects than their low-latitude counterparts.

While both species became infected in our experiment, in *Ra* IL was lower and *Bd*-mediated mortality was only a fraction of the mortality experienced by *Bb*. These results agree with previous studies showing that brown frogs, such as *Ra* have higher tolerance to *Bd*, while *Bb*, like many other bufonids, is more susceptible to *Bd*-infection (Bosch & Martínez-Solano 2006, Garner et al. 2011, Gahl et al. 2012, Balaz et al. 2014, Bielby et al. 2015). When comparing susceptibility to infection and *Bd*-mediated mortality between two anuran species, Bielby et al. (2015) found that *R. temporaria*, closely related to *Ra*, was resistant to infection even at high doses, while *Bb* showed near complete infection and dose-dependent mortality. Since *Ra* has higher infection prevalence in in the wild (Meurling et al. 2020) and higher infection tolerance (this study), we suggest that *Ra* may act as a reservoir species and a possible vector for *Bd*-transmission to more sensitive species such as *Bb*. Indeed, Kärvemo et al. (2019) showed that *Bb* populations coexisting with *Ra* had higher *Bd*-prevalence than populations breeding in ponds without *Ra*.

We found a clear difference in survival between northern and southern populations especially in *Bb*. Northern individuals were smaller at the time of infection than southern individuals, and our analyses suggest that the survival difference was mainly mediated by body size. This is in accordance with previous studies (Bradley et al. 2015, Burrow et al. 2017) showing that smaller individuals were more vulnerable to *Bd* infection. Smaller individuals may have less developed immune system which may render them more vulnerable to disease (Møller et al. 1998, Burrow et al. 2017). Smaller individuals may also be more vulnerable to *Bd*-mediated water loss as they have larger surface area to body mass ratio. Increased water loss via sloughing is an important symptom in chytridiomycosis, which may render smaller individuals more sensitive to *Bd* infection (Russo et al. 2018, Wu et al. 2019).

Our results suggest that much of the differences in *Bd*-mediated mortality can be explained by size differences between the regions. As we raised the tadpoles under common garden conditions, the differences in body mass most likely have a genetic origin. Two additional, not mutually exclusive, explanations may further explain higher mortality in the northern populations. Firstly, northern populations may have less effective immune systems because of reduced genetic variation due to postglacial colonization processes (Hewitt 2000, see Cortazar-Chinarro et al. 2017, Rödin-Mörch et al. 2019 for *Ra*, Thörn et al. 2021 for *Bb*), or lower pathogen abundance at higher latitudes (Schemske et al. 2009). This hypothesis gains support from the fact that MHC variation in both our study species is lower at higher latitudes (Cortázar-Chinarro et al. 2017, Meurling 2019). Moreover, *Bd*-mediated survival in *Bb* seems to be linked with certain MHC alleles (Meurling 2019), as also found in other species (Savage & Zamudio 2011, Savage et al. 2018, Kosch et al. 2019). Secondly, higher larval development rates in the northern populations may trade off with disease resistance (Johnson et al. 2011, Woodhams et al. 2016). Also this hypothesis is indirectly supported by the facts that more time-constrained populations have higher development rates in both our study species (Luquet et al. 2015, 2019) and that *Ra* tadpoles experimentally induced to develop faster have weaker immune response (Murillo-Rincon et al. 2017). Additional studies focusing on *Bd* resistance in known MHC and developmental genotypes would be highly interesting.

*Bd* infection had clear negative effects on growth in both species. As body size is positively related to fitness in juvenile amphibians (Earl and Whiteman 2015), these results suggest that *Bd* may have sublethal fitness effects. For example, hibernation success is often positively related to body size and failing to reach a sufficient size before hibernation can greatly reduce overwinter survival (Altwegg and Reyer 2003). This can be especially detrimental at higher latitudes where the hibernation period can reach eight months. Small body size may also lead to higher risk of predation, delayed maturation and lower ability to compete for resources and mates (reviewed in Earl and Whiteman 2015). In the long run, these effects may decrease population growth rate and ability to cope with environmental changes such as higher temperature due to climate change. In our case, even if survival of *Ra* was not strongly affected by *Bd* infection, the results suggest that sublethal effects of infection mediated by body size may still lower individual and population fitness.

We found relatively little evidence for differences in pathogenicity between the two *Bd* isolates. However, we found significant treatment × size interaction in survival of *Bb* where survival was more strongly size-dependent when toadlets were infected with the UK strain. These results suggest that individuals infected with the UK strain may relatively quickly reach a size where the lethality of *Bd* is reduced, while *Bd*-mediated mortality induced by the Swedish strain is less size-dependent. This is especially the case in southern individuals which are larger at metamorphosis, while the smaller northern individuals stay longer in the vulnerable size classes.

A potential caveat in our study is that as we used laboratory-raised (but wild-collected) individuals which may not have developed as diverse community of skin microbiota as wild individuals. Indeed, captive amphibians often have a reduced and less varied bacterial community than wild populations of the same species (Antwis et al. 2014, Bataille et al. 2016). As skin microbiome plays an important role in defending against fungal and other pathogens, this could impact the ability of amphibians reared in captivity to respond to *Bd* infection (Harris et al. 2009, Walke et al. 2015, Madison et al. 2017, Woodhams et al. 2018). We currently lack knowledge on the skin microbiomes of our study species and if these differ between geographical regions. Microbiome studies are needed for additional insight on factors behind the high mortality found in this study.

*Bd* is widespread in the southern parts of Sweden (Kärvemo et al. 2018, 2019, Meurling et al. 2020). However, in a pattern similar to much of Europe (Lips 2016, Scheele et al. 2019), no cases of chytridiomycosis or unusual die-offs have been found in Sweden. Our experimental results suggest that even though no negative effects of the infection have been seen in the wild, this might not be the complete picture. It is currently unclear how well the present results translate to natural conditions, but we note that *Bd* causes sublethal effects in terms of reduced movements and body condition in wild Scandinavian amphibians (Kärvemo et al. 2019, 2020). Furthermore, the lethality of *Bd* is highly dependent on environmental conditions, including temperature (e.g., Novakowski et al. 2106, Mosher et al. 2018, Cohen et al. 2019), and relatively minor elevations in mortality may risk long-term survival of *Bd*-infected amphibian populations (Muths et al. 2011; Spitzen-van der Sluijs et al. 2017). Two important conclusions can be drawn. Firstly, very few surveys have been conducted in northern Scandinavia (Meurling et al. 2020). As populations at higher latitudes can be more vulnerable to infection, it is important to investigate the occurrence of *Bd* in these areas and, if still possible, prevent or limit the northward spread of the fungus. Secondly, we showed that infection leads to higher mortality and reduced body size. These, in turn, can lead to reduced population growth rates in the long-term even in the absence of major mortality effects. As the potential negative effects of *Bd* on population growth can be relatively subtle and difficult to detect (Doddington et al. 2013, Spitzen-van der Sluijs et al. 2017, Mosher et al. 2018), long-term monitoring of amphibian populations is of high importance.

## Acknowledgements

We thank Lola Brooks and Trent Garner for discussions and providing the *Bd*-strains.

## Declarations

### Ethics approval

The study was conducted with a permit (C28/15) from Uppsala ethical committee for animal experiments and collection permits from the county administrative boards in Skåne and Norrbotten.

### Availability of data and materials

The data will be deposited in DRYAD upon acceptance.

### Authors’ contributions

SM, MCC, JH and AL conceived and designed the experiments, SM, MCC and DÅ performed the experiments and EÅ provided advice and logistic help, SM and MLS analysed the data, SM and AL wrote the paper with input from all the authors.

---

**Table S1.** Coordinates for the collection sites.

| Species | Region | Population | N | E |
| --- | --- | --- | --- | --- |
| *Rana arvalis* | North | NG1 | 65.519974 | 21.685978 |
|  |  | NG2 | 65.488998 | 21.378151 |
|  | South | M | 55.699774 | 13.360416 |
|  |  | K | 55.722114 | 13.284693 |
| *Bufo bufo* | North | NP1 | 65.583139 | 22.319458 |
|  |  | NP2 | 65.56554 | 22.37404 |
|  | South | PM | 56.217897 | 13.731548 |
|  |  | PH | 55.539031 | 14.009702 |

**Table S2.** Results from final general linear models on infection load. a) *R. arvalis*, b) *B. bufo.*

|  | Sum of squares | Df | *F* | *P* |
| --- | --- | --- | --- | --- |
| Size | 5.5 | 1 | 7.54 | **0.008** |
| Region | 8.8 | 1 | 12.11 | **< 0.001** |
| Size x Region | 4.7 | 1 | 6.50 | **0.013** |
| Residuals | 50.3 | 69 |  |  |

b)
|  | Sum of squares | Df | *F* | *P* |
| --- | --- | --- | --- | --- |
| Bd-strain | 0.8 | 1 | 1.70 | 0.196 |
| Size | 31.9 | 1 | 64.45 | **< 0.001** |
| Strain x Size | 1.8 | 1 | 3.66 | 0.059 |
| Residuals | 43.0 | 87 |  |  |

**Table S3.**
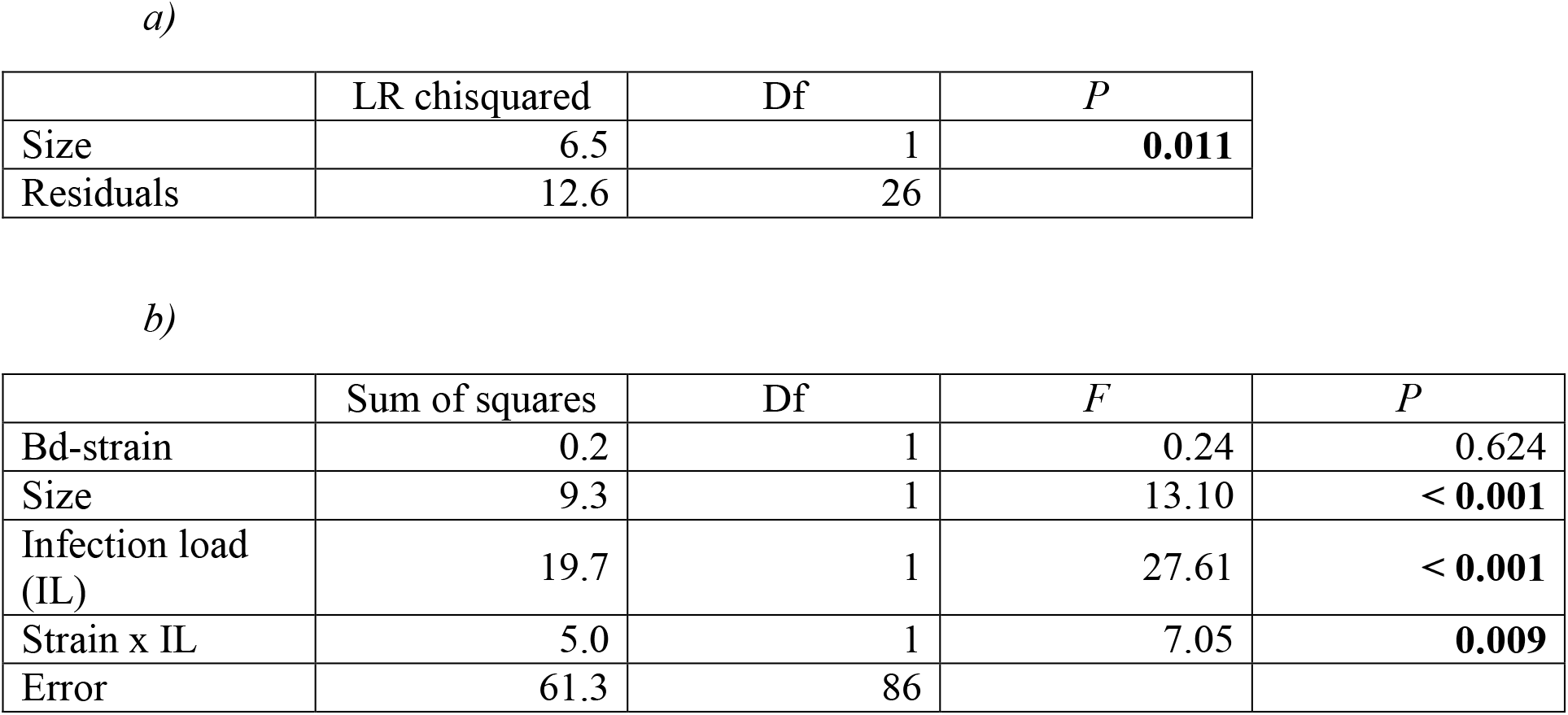
Results from final generalized linear models on survival. a) *R. arvalis*: the analyses only cover the northern population and the two *Bd*-treatments as survival in the southern population and control treatment were complete in this species. b) *B. bufo:* only the two *Bd*-treatments are included as survival was complete in the control treatment in this species.

|  | LR chisquared | Df | *P* |
| --- | --- | --- | --- |
| Size | 6.5 | 1 | **0.011** |
| Residuals | 12.6 | 26 |  |

b)
|  | Sum of squares | Df | *F* | *P* |
| --- | --- | --- | --- | --- |
| Bd-strain | 0.2 | 1 | 0.24 | 0.624 |
| Size | 9.3 | 1 | 13.10 | **< 0.001** |
| Infection load (IL) | 19.7 | 1 | 27.61 | **< 0.001** |
| Strain x IL | 5.0 | 1 | 7.05 | **0.009** |
| Error | 61.3 | 86 |  |  |

**Table S4.** Results from general linear models on growth. a*) R. arvalis*, b) *B. bufo* all individuals, c) *B. bufo*, *Bd*-infected individuals only

|  | Sum of squares | Df | *F* | *P* |
| --- | --- | --- | --- | --- |
| *Bd* strain | 0.000131 | 2 | 3.13 | **0.048** |
| Region | 0.000001 | 1 | 0.05 | 0.827 |
| Size | 0.000143 | 1 | 6.84 | **0.010** |
| Infection load (IL) | 0.000129 | 1 | 6.16 | **0.015** |
| Region x Size | 0.000053 | 1 | 2.54 | 0.114 |
| Size x IL | 0.000150 | 1 | 7.21 | **0.008** |
| Residuals | 0.0002233 | 107 |  |  |

b)
|  | Sum of squares | Df | *F* | *P* |
| --- | --- | --- | --- | --- |
| *Bd*-strain | 0.000209 | 2 | 2.43 | 0.092 |
| Region | 0.000117 | 1 | 2.72 | 0.101 |
| Size | 0.000006 | 1 | 0.14 | 0.712 |
| Infection load (IL) | 0.000336 | 1 | 7.82 | **0.006** |
| Strain x Size | 0.000252 | 2 | 2.94 | 0.056 |
| Region x Size | 0.000937 | 1 | 3.06 | 0.083 |
| Region x IL | 0.000131 | 1 | 3.66 | 0.059 |
| Size x IL | 0.000391 | 1 | 21.84 | **< 0.001** |
| Residuals | 0.005667 | 132 |  |  |

c)
|  | Sum of squares | Df | *F* | *P* |
| --- | --- | --- | --- | --- |
| *Bd*-strain | 0.000003 | 1 | 0.06 | 0.812 |
| Region | 0.000001 | 1 | 0.02 | 0.877 |
| Size | 0.000494 | 1 | 10.67 | **0.002** |
| Infection load (IL) | 0.000513 | 1 | 11.09 | **0.001** |
| Strain x Region | 0.000102 | 1 | 2.20 | 0.142 |
| Region x Size | 0.000814 | 1 | 17.61 | **< 0.001** |
| Size x IL | 0.000416 | 1 | 9.00 | **0.004** |
| Residuals | 0.003838 | 83 |  |  |

**Fig. S1.**
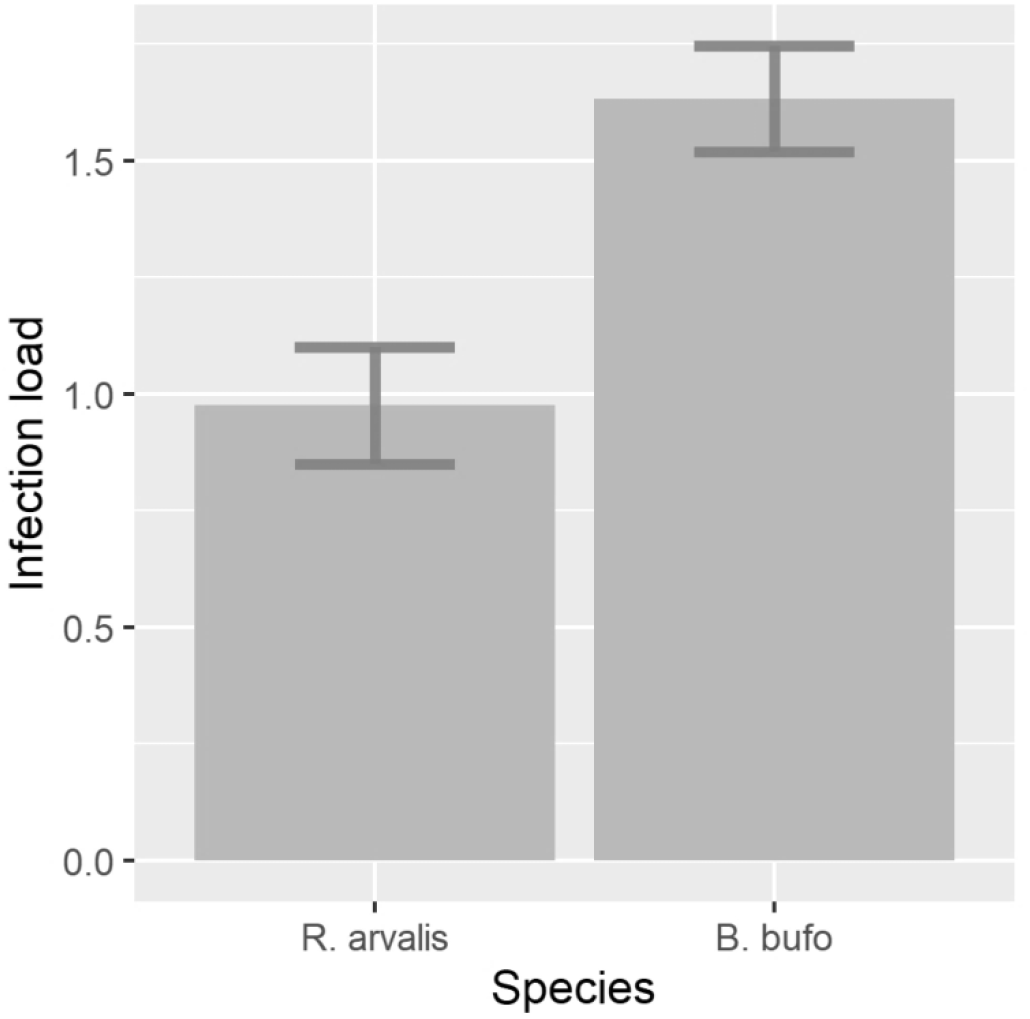
Infection load (genomic equivalents) in *R.arvalis* and *B.bufo*.

## References

Altwegg R, Reyer HU (2003) Patterns of natural selection on size at metamorphosis in water frogs. Evolution 57:872–882

Antwis RE, Haworth RL, Engelmoer DJ et al.(2014) Ex situ diet influences the bacterial community associated with the skin of red-eyed tree frogs (*Agalychnis callidryas*). PLoS One 9:e85563

APHA. 1985. Standard methods for the examination of water and wastewater. 16th ed. American Public Health Association, Washington, DC.

Balaz V, Voros J, Civis P et al. (2014) Assessing risk and guidance on monitoring of *Batrachochytrium dendrobatidis* in Europe through identification of taxonomic selectivity of infection. Conserv Biol 28:213–223

Bataille A, Fong JJ, Cha M et al. (2013) Genetic evidence for a high diversity and wide distribution of endemic strains of the pathogenic chytrid fungus *Batrachochytrium dendrobatidis* in wild Asian amphibians. Mol Ecol 22:4196–4209.

Bataille A, Lee-Cruz L, Tripathi B et al. (2016) Microbiome variation across amphibian skin regions: Implications for chytridiomycosis mitigation efforts. Microb Ecol 71:221–232

Berger L, Speare R, Daszak P et al. (1998) Chytridiomycosis causes amphibian mortality associated with population declines in the rain forests of Australia and Central America. Proc Natl Acad Sci USA 95:9031–9036

Becker CG, Bletz MC, Greenspan SE et al. (2019) Low-load pathogen spillover predicts shifts in skin microbiome and survival of a terrestrial-breeding amphibian. Proc R Soc B 286:20191114

Bielby J, Fisher MC, Clare FC et al. (2015) Host species vary in infection probability, sub-lethal effects, and costs of immune response when exposed to an amphibian parasite. Sci Rep 5:10828

Bosch J, Martínez-Solano I (2006) Chytrid fungus infection related to unusual mortalities of *Salamandra salamandra* and *Bufo bufo* in the Peñalara Natural Park, Spain. Oryx 40:84–89

Boyle DG, Boyle DB, Olsen V et al. (2004) Rapid quantitative detection of chytridiomycosis (*Batrachochytrium dendrobatidis*) in amphibian samples using real-time Taqman PCR assay. Dis Aquat Org 60:141–148

Bradley PW, Gervasi SS, Hua J et al. (2015) Differences in sensitivity to the fungal pathogen *Batrachochytrium dendrobatidis* among amphibian populations. Conserv Biol 29:134–1356

Burrow AK, Rumschlag SL, Boone MD (2017) Host size influences the effects of four isolates of an amphibian chytrid fungus. Ecol Evol 7:9196–9202

Casadevall A (2007) Determinants of virulence in the pathogenic fungi. Fungal Biol Rev 21:130–132

Campbell L, Bower DS, Clulow S et al. (2019) Interaction between temperature and sublethal infection with the amphibian chytrid fungus impacts a susceptible frog species. Sci Rep 9:83

Cohen JM, McMahon TA, Ramsay C et al. (2019) Impacts of thermal mismatches on chytrid fungus *Batrachochytrium dendrobatidis* prevalence are moderated by life stage, body size, elevation and latitude. Ecol Lett 22:817–825

Cortázar-Chinarro M, Lattenkamp EZ, Meyer-Lucht Y et al. (2017) Drift, selection, or migration? Processes affecting genetic differentiation and variation along a latitudinal gradient in an amphibian. BMC Evol Biol 17:189

Dang TD, Searle CL, Blaustein AR (2017) Virulence variation among strains of the emerging infectious fungus *Batrachochytrium dendrobatidis* in multiple amphibian host species. Dis Aquat Org 124:233–239

Daszak P, Cunningham AA, Hyatt AD (2000) Emerging infectious diseases of wildlife-- Threats to biodiversity and human health. Science 287:443–449

Doddington BJ, Bosch J, Oliver JA et al. (2013) Context-dependent amphibian host population response to an invading pathogen. Ecology 94:1795–1804

Earl JE, Whiteman HH (2015) Are commonly used fitness predictors aAccurate? A meta-analysis of amphibian size and age at metamorphosis. Copeia 103:297–309

Ebert D (2008) Host-parasite coevolution. insights from the *Daphnia*-parasite model system. Curr Opin Microbiol 11:290–301

Ellison AR, Tunstall T, DiRenzo GV et al. (2014) More than skin deep: functional genomic basis for resistance to amphibian chytridiomycosis. Genome Biol Evol 7:286–298

Farrer RA, Weinert LA, Bielby J et al. (2011) Multiple emergences of genetically diverse amphibian-infecting chytrids include a globalized hypervirulent recombinant lineage. Proc Natl Acad Sci USA 108:18732–18736

Fisher MC, Garner TW, Walker SF (2009) Global emergence of *Batrachochytrium dendrobatidis* and amphibian chytridiomycosis in space, time, and host. Annu Rev Microbiol 63:291–310

Fisher MC, Henk DA, Briggs CJ et al. (2012) Emerging fungal threats to animal, plant and ecosystem health. Nature 484:186

Gahl MK, Longcore JE, Houlahan JE (2012) Varying responses of rortheastern North American amphibians to the chytrid pathogen *Batrachochytrium dendrobatidis*. Conserv Biol 26:135–141

Garner TWJ, Rowcliffe JM, Fisher MC (2011) Climate change, chytridiomycosis or condition: an experimental test of amphibian survival. Glob Change Biol 17:667–675

Gervasi S, Foufopoulos J (2007) Costs of plasticity: Responses to desiccation decrease post-metamorphic immune function in a pond-breeding amphibian. Funct Ecol 22:100–108

Gosner KL (1960) A Simplified table for staging anuran embryos and larvae with notes on identification. Herpetologica 16:183–190

Greenspan SE, Lambertini C, Carvalo T et al. (2018) Hybrids of amphian chytrid show high virulence in native hosts. Sci Rep 8:9600

Rowley-Harris RN, Brucker RM, Walke JB et al. (2009) Skin microbes on frogs prevent morbidity and mortality caused by a lethal skin fungus. ISME J 3:818–824

Herczeg D, Ujszegi J, Kasler A et al. 2021. Host-multiparasite interactions in amphibians: a review. Parasit Vectors 14:296

Hewitt G (2000) The genetic legacy of the Quaternary ice ages. Nature 405:907–913

Hyatt AD, Boyle DG, Olsen V et al. (2007) Diagnostic assays and sampling protocols for the detection of *Batrachochytrium dendrobatidis*. Dis Aquat Org 73:175–192

Johnson PTJ, Kellermanns E, Bowerman J (2011) Critical windows of disease risk: amphibian pathology driven by developmental changes in host resistance and tolerance. Funct Ecol 25:726–734

Kärvemo S, Laurila A, Höglund J (2019) Urban environment and reservoir host species are associated with *Batrachochytrium dendrobatidis* infection prevalence in the common toad. Dis Aquat Org 134:33–42

Kärvemo S, Meurling S, Berger D et al. (2018) Effects of host species and environmental factors on the prevalence of *Batrachochytrium dendrobatidis* in northern Europe. PLoS ONE 13:e0199852

Kärvemo S, Wikström G, Widenfalk LA et al. (2020) Chytrid fungus dynamics and infections associated with movement distances in a red-listed amphibian. J Zool 311:164–174

Kosch TA, Silva CNS, Brannelly LA et al. (2019) Genetic potential for disease resistance in critically endangered amphibians decimated by chytridiomycosis. Anim Conserv 22:238–250

Laine A-L, Burdon JJ, Dodds PN et al. (2011) Spatial variation in disease resistance: from molecuales to metapopulations. J Ecol 99:96–112.

Lips KR (2016) Overview of chytrid emergence and impacts on amphibians. Phil Trans R Soc B 371:20150465

Lorch JM, Knowles S, Lankton JS et al. (2016) Snake fungal disease: an emerging threat to wild snakes. Phil Trans R Soc B 371:20150457

Luquet E, Léna JP, Miaud C et al. (2015) Phenotypic divergence of the common toad (*Bufo bufo*) along an altitudinal gradient: evidence for local adaptation. Heredity 114:69–79

Luquet E, Rödin Mörch P, Cortázar-Chinarro M et al. (2019) Post-glacial colonization routes coincide with a life-history breakpoint along a latitudinal gradient. J Evol Biol 32:356–368

Martin-Torrijos L, Campos Llach M, Pou_Rovira Q et al. (2017) Resistance to crayfish plague, *Aphanomyces astaci* (Oomycota) in the endangered freshwater crayfish species *Austropotamobius pallipes*. PLoS ONE 12:e0181226

Madison JD, Berg EA, Abarca JG et al. (2017) Characterization of *Batrachochytrium dendrobatidis* inhibiting bacteria from amphibian populations in Costa Rica. Front Microbiol 8:290

Meurling S (2019) The response in native wildlife to an invading pathogen: Swedish amphibians and *Batrachochytrium dendrobatidis*. PhD thesis, Uppsala University.

Meurling S, Kärvemo S, Chondrelli N et al. (2020) Occurrence of *Batrachochytrium dendrobatidis i*n Sweden: higher infection prevalence in southern species. Dis Aquat Org 140:209–218

More S, Angel Miranda M, Bicout D et al.(2018) Risk of survival, establishment and spread of Batrachochytrium salamandrivorans (*Bsal*) in the EU. EFSA J 16:e05259

Mlller AP, Christe P, Erritzoe J et al. (1998). Condition, disease and immune defence. Oikos 83:301o306

Mosher BA, Bailey LL, Muths E et al. (2018) Host-pathogen metapopualtion suggest high elevation refugia for boreal toads. Ecol Appl 28:926–937

Murillo-Rincon AP, Laurila A, Orizaola G (2017) Compensating for delayed hatching reduces offspring immune response and increases life-history costs. Oikos 126:565–571

Muths E, Scherer RD, Pilliod DS (2011) Compensatroy effects of recruitment and survival when amphibians are perturbed by disease. Ecol Appl 48:873–879

Nowakowski AJ, Whitfield SM, Eskew EA et al. (2016) Infection risk decreases with increasing mismatch in host and pathogen environmental tolerances. Ecol Lett 19:1051–1061

O’Hanlon SJ, Rieux A, Farrer RA et al. (2018) Recent Asian origin of chytrid fungi causing global amphibian declines. Science 360:621–627

Olson DH, Aanensen DM, Ronnenberg KL et al. (2013) Mapping the global emergence of *Batrachochytrium dendrobatidis*, the amphibian chytrid fungus. PLoS ONE 8:e56802

Palo JU, O’Hara RB, Laugen AT et al. (2003) Latitudinal divergence of common frog *(Rana temporaria*) life history traits by natural selection: evidence from a comparison of molecular and quantitative genetic data. Mol Ecol 12:1963–1978

Pennisi E (2010) Armed and dangerous. Science 327:804–805

R Core Team (2018) R: A Language and Environment for Statistical Computing. R Foundation for Statistical Computing, Vienna, Austria

Richards-Zawacki CL (2010) Thermoregulatory behaviour affects prevalence of chytrid fungal infection in a wild population of Panamanian golden frogs. Proc R Soc B 277:519–528

Rödin-Mörch P, Luquet E, Meyer-Lucht Yet al. (2019) Latitudinal divergence in a widespread amphibian: Contrasting patterns of neutral and adpative genomic variation. Mol Ecol 28:2996–3011

Russo CJM, Ohmer MEB, Cramp RL et al. (2018) A pathogenic skin fungus and sloughing exacerbate cutaneous water loss in amphibians. J Exp Biol 221

Savage AE, Zamudio KR (2011) MHC genotypes associate with resistance to a frog-killing fungus. Proc Natl Acad Sci USA 108:16705–16710

Savage AE, Mulder KP, Torret T et al. (2018) Lost but nmot forgotten: MHC genotypes predict overwinter survival despite depauparate MHC diversity in a declining frog. Conserv Genet 19:309–322

Scheele BC, Hunter DA, Brannelly LA et al. (2017) Reservoir-host amplification of disease impact in an endangered amphibian. Conserv Biol 31:592–600

Scheele BC, Pasmans F, Skerratt LF et al. (2019) Amphibian fungal panzootic causes catastrophic and ongoing loss of biodiversity. Science 363:1459–1463

Schemske DW, Mittelbach GG, Cornell HV et al. (2009) Is there a latitudinal gradient in the importance of biotic interactions? Annu Rev Ecol Evol Syst 40:245–269

Sillero N, Campos J, Bonardi A et al. (2014) Updated distribution and biogeography of amphibians and reptiles of Europe. Amph-Reptil 35:1–31

Skerratt LF, Berger L, Speare R et al. (2007) Spread of chytridiomycosis has caused the rapid global decline and extinction of frogs. EcoHealth 4:125–134

Spitzen - van der Sluijs A, Canessa S, Martel A 2017 Fragile coexistence of a global chytrid pathogen with amphibian populations is medaited by environment and demography. Proc R Soc B 284:20171444

Thörn F, Rödin-Mörch P, Cortazar-Chinarro M et al. (2021) The effects of drift and selection on latitudinal genetic variation in Scandinavian common toads (*Bufo bufo*) following postglacial recolonization. Heredity 126:656–667

Voyles J, Woodhams DC, Saenz V et al. (2018) Shifts in disease dynamics in a tropical amphibian assemblage are not due to pathogen attenuation. Science 359:1517

Walke JB, Becker MH, Loftus SC et al. (2015) Community structure and function of amphibian skin microbes: an experiment with bullfrogs exposed to a chytrid fungus. PLoS ONE 10:e0139848

Wang SP, Liu CH, Wilson AB et al. (2017) Pathogen richness and abundance predict patterns of adaptive major histocompatibility complex variation in insular amphibians. Mol Ecol 26:4671–4685

Wibbelt G, Kurth A, Hellmann D et al. (2010) White-nose syndrome fungus (*Geomyces destructans*) in bats, Europe. Emerg Infect Dis 16:1237–1243

Woodhams DC, Bletz M, Kueneman J et al. (2016) Managing amphibian disease with skin microbiota. Trends Microbiol 24:161–164

Woodhams DC, LaBumbard BC, Barnhart KL et al. (2018) Prodigiosin, violacein, and volatile organic compounds produced by widespread cutaneous bacteria of amphibians can inhibit two *Batrachochytrium* fungal pathogens. Microb Ecol 75:1049–1062

Wu NC, McKercher C, Cramp RL et al. (2019) Mechanistic basis for loss of water balance in green tree frogs infected with a fungal pathogen. Am J Physiol Regul Integr Comp Physiol 317:R301–R311

Zeisset I, Beebee TJC (2014) Drift rather than selection dominates MHC Class II Allelic diversity patterns at the biogeographical range scale in natterjack toads *Bufo calamita*. PLoS ONE 9:e100176

